# Isolation and genome characterization of Lloviu virus from Italian Schreibers’ bent-winged bats

**DOI:** 10.1101/2022.12.05.519067

**Authors:** Gábor E. Tóth, Adam J. Hume, Ellen L. Suder, Safia Zeghbib, Ágota Ábrahám, Zsófia Lanszki, Zsaklin Varga, Zsófia Tauber, Fanni Földes, Brigitta Zana, Dino Scaravelli, Maria Teresa Scicluna, Andrea Pereswiet-Soltan, Tamás Görföl, Calogero Terregino, Paola De Benedictis, Isabel Garcıa-Dorival, Covadonga Alonso, Ferenc Jakab, Elke Mühlberger, Stefania Leopardi, Gábor Kemenesi

## Abstract

Lloviu cuevavirus (LLOV) was the first identified member of *Filoviridae* family outside the *Ebola* and *Marburgvirus* genera. A massive die-off of Schreibers’ bent-winged bats (*Miniopterus schreibersii*) in the Iberian Peninsula in 2002 led to its discovery.

Studies with recombinant and wild-type LLOV isolates confirmed the susceptibility of human-derived cell lines and primary human macrophages to LLOV infection *in vitro*. Based on these data, LLOV is now considered as a potential zoonotic virus with unknown pathogenicity to humans and bats.

We examined bat samples from Italy for the presence of LLOV in an area outside of the currently known distribution range of the virus. We detected one positive sample from 2020, sequenced the complete coding sequence of the viral genome and established an infectious isolate of the virus. In addition, we performed the first comprehensive evolutionary analysis of the virus, using the Spanish, Hungarian and the Italian sequences.

The most important achievement of this article is the establishment of an additional infectious LLOV isolate from a bat sample using the SuBK12-08 cells, demonstrating that this cell line is highly susceptible to LLOV infection. These results further confirms the role of these bats as the host of this virus, possibly throughout their entire geographic range. This is an important result to further understand the role of bats as the natural hosts for zoonotic filoviruses.

## Study

Lloviu cuevavirus (LLOV) was the first identified member of Filoviridae family outside the Ebola and Marburgvirus genera. A massive die-off of Schreibers’ bent-winged bats (*Miniopterus schreibersii*) in the Iberian Peninsula in 2002 led to its discovery. Almost complete genome sequence of the virus was retrieved with various sequencing approaches, although the infectious virus was not isolated from these bats at that time [1]. It has been reported that Schreibers’ bent-winged bats in Spain and Hungary are LLOV-seropositive, confirming that the virus circulates there [2,3]. Viral RNA was sequenced from Spanish and Hungarian samples, and recently, an infectious isolate was established from the blood sample of a naturally infected bat in Hungary [1,3,4].

Studies with recombinant and wild-type LLOV isolates confirmed the susceptibility of human-derived cell lines and primary human macrophages to LLOV infection *in vitro*. Based on these data, LLOV is now considered as a potential zoonotic virus with unknown pathogenicity to humans and bats [3,5].

The ability of LLOV infection to cause disease in bats is unclear. Although no causative relationship has been established between Schreibers’ bent-winged bat die-off events and circulation of LLOV within bat populations to date, the correlation is noteworthy [3,6]. The recent isolation of the virus from Schreibers’ bent-winged bats in Hungary confirmed the role of these bats as hosts for LLOV and perhaps as a reservoir species. It is unclear if the first detection of the virus in Spain coincided with an initial introduction of LLOV to European *M. schreibersii* populations. Alternatively, it is possible that LLOV was already broadly circulating in these bats across the continent.

To further explore the occurrence of LLOV in the wider distribution range of these bats and better understand its ecology and evolution, we screened 282 samples from Schreibers’ bent-winged bats from four different locations in northern and central Italy between 2018 and 2021 (Figure 1A), including 173 blood clots and 108 lung sections. Lung samples were obtained in the framework of the passive surveillance of bats for rabies from necropsies of dead adult and neonate bats. Archived samples, collected since 2018, represented samples collected throughout the whole year, including in the winter months. Blood samples were obtained during four active surveillance campaigns between late summer 2020 (August, September, October) and spring 2021 (April), in order to avoid disturbing the bat colonies during hibernation and the first month after birth. Bats were captured during the day with hand-nets in full compliance with best practice and national and European regulations, in derogation to the Council Directive 92/43/EEC of 21 May 1992 on the conservation of natural habitats and of wild fauna and flora (authorization 38025 of 13/08/2020 and 6831 of 15/02/2021 released by the Italian Ministry for the Ecological Transition).

**Figure 1.**
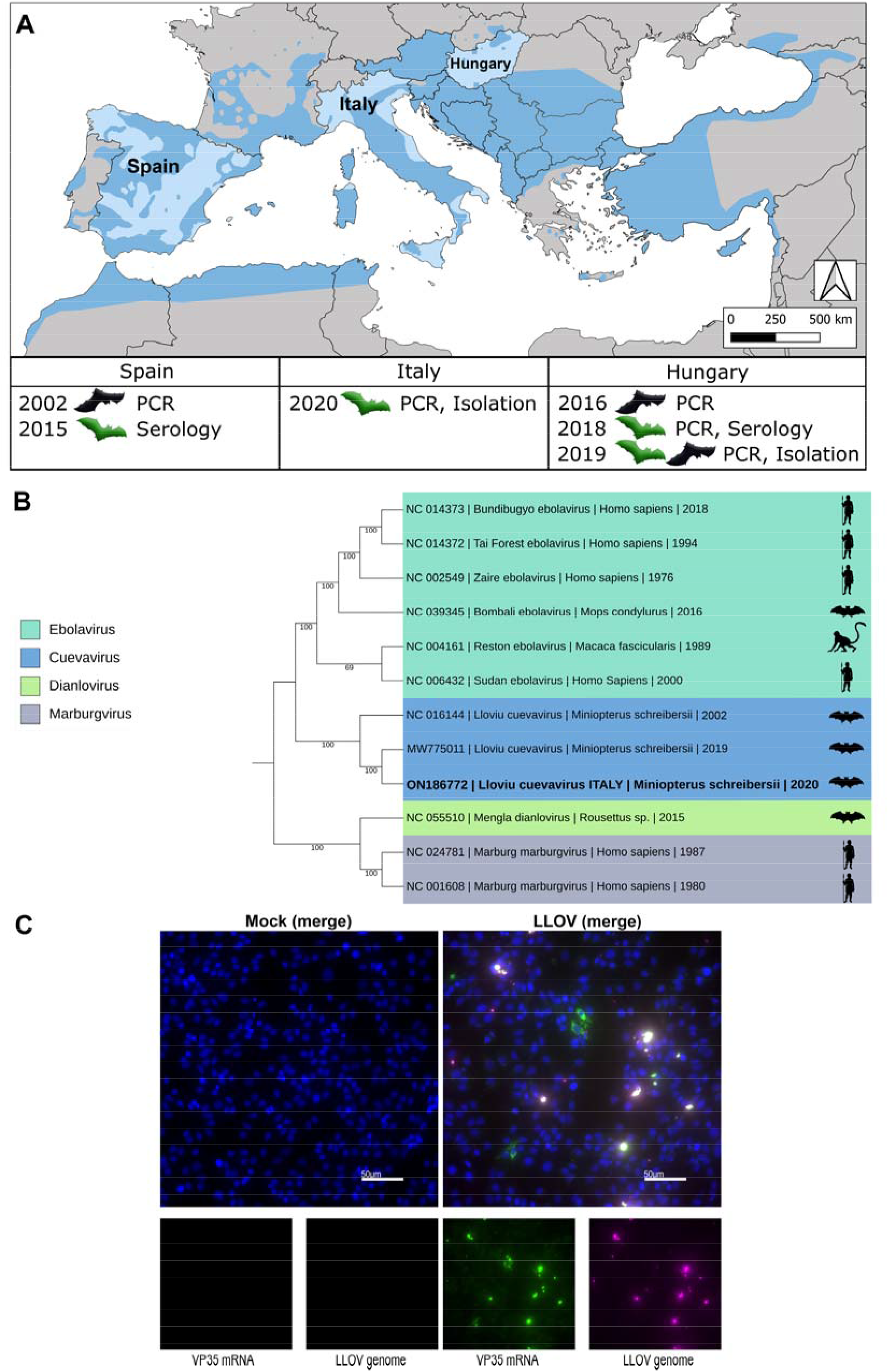
(A) Geographic distribution of Schreibers’ bats and identified locations of LLOV detection. The map shows the currently known distribution range of Schreibers’ bats (darker blue). Countries in which LLOV has been found are named on the map. The year and methods of detection are indicated below the countries of sample origin. Bat pictograms in normal position (green) indicate samples from live animals while upside down bats (black) indicate samples from bat carcasses. (B) Phylogenetic analyses of mammalian-associated filovirus reference genomes, including the available bat-derived LLOV sequences. The Maximum Likelihood tree was built using general time reversible model of substitution with gamma distributed rate variation and was tested with 1000 bootstrap replicates. (C) SuBK12-08 cells were left uninfected (mock) or infected with the Italian LLOV isolate. At 1-day post-infection, cells were fixed with 10% formalin and stained by RNA FISH using probes targeting the negative sense viral genome (magenta) and the positive sense VP35 mRNA (green). Cell nuclei are stained with DAPI (blue).

RNA was extracted from samples using Direct-Zol RNA Miniprep kit (Zymo Research) according to the manufacturer’s recommendations. We performed LLOV-specific real-time RT-PCR screening, targeted viral genome sequencing and evolutionary analysis, complemented with *in vitro* isolation experiments as described before [3]. For LLOV RNA detection, a novel TaqMan PCR assay was developed and optimized using the Lloviu cuevavirus isolate Hungary/2019/378 (GenBank: MZ541881.1). We used the following oligonucleotide sequences: LLOV-Fw-scr1: 5’-AAGCATTTCCGAGTAATATGATGGTTG-3’, LLOV-Rev-scr1: 5’-TACATGGTCTCCTAGATTGCCCTG-3’, LLOV-Prob-scr1: 5’-FAM-CCTGATGAAGGAGAGTTTCTTTCTG-ZEN-3’ with the qRT-PCR Brilliant III Probe Master Mix (Agilent Technologies) under these conditions: 50°C for 10 minutes, 95°C for 3 minutes and 50 cycles of 95°C for 5 seconds, 60°C for 30 seconds.

While all 108 lung samples tested negative for LLOV RNA, one of the 173 blood clot samples was positive with a Ct value of 21.07, corresponding to approximately 3.8x10^8^ genomic copies per mL. The sample belonged to a set of 75 blood clots collected in September 2020 from a single colony. Amplicon-based, targeted Nanopore sequencing was used to determine the complete coding sequence of the viral genome. Combined, 615,045 reads were mapped to the reference genome which resulted in a 27,489.5x mean coverage (Figure S1A). The retrieved sequence was 18,861 nucleotides in length, covering all seven viral genes (NP, VP35, VP40, GP, VP30, VP24, and L) (Genbank: ON186772). Compared to the reference LLOV genome from Spain (Genbank: NC016144.1) and the Hungarian LLOV sequence (GenBank: MW775011), the sequence from the Italian LLOV sample showed 99.13% and 99.86% identity at nucleotide level, respectively. The phylogenetic analysis showed that the Italian LLOV isolate is more closely related to the Hungarian isolate (Figure 1B).

Analysis of the mutational landscape gives the first insight into the evolution of Lloviu cuevavirus. Comparing to the published Spanish LLOV sequence, the dN/dS ratios of each gene indicate that only GP (Hungarian: 1.38, Italian: 1.5) had a ratio above 1, implying the presence of positive selection pressure. The highest rate of synonymous mutations were found in the VP24 gene. Using the Spanish LLOV sequence as a baseline, we observed a roughly linear mutational rate for the Hungarian and Italian LLOV sequences with respect to the time of sample collection (Figure S1B). Non-synonymous mutations are summarized in Table S1. While several amino acid coding changes were found throughout the LLOV genome most of these were localized in GP (Table S1). Interestingly, some of these mutations have been previously reported and linked to virus adaptation into a new species, such is the case of amino acid positions 465 and 493 that have been found in EBOV GP when adapted to a new host [7]. Another interesting coding change we identified was in amino acid position 16 (F16L) of VP24. A similar but inverted change was found in aa position 26 (L26F) for EBOV VP24. This mutation was described as a molecular determinant of virulence in guinea pig model, with this single amino acid change in EBOV proving to be sufficient to cause increased pathogenesis and death in guinea pigs [8]. This observation was further confirmed in another study where EBOV amino acid variations associated with increased pathogenesis were studied [7]. Considering the phylogenetic relatedness and functional similarities of LLOV to EBOV, these mutations are prominent targets of functional experiments in the future to better understand LLOV evolution and pathogenesis.

In addition to sequence-based analyses we conducted *in vitro* isolation experiments using the LLOV-positive sample to infect the *Miniopterus*-derived cell line SuBK12-08 [9]. After the first passage of the supernatant, a massive cytopathic effect (CPE) was observed in the cells, similar to previous LLOV isolation studies [3]. To verify the presence of replicative LLOV, we performed RNA fluorescent in situ hybridization (FISH) staining with LLOV RNA-specific probes.. We observed accumulation of LLOV genomic RNA in virus-induced inclusion bodies and a diffuse distribution of VP35 mRNA throughout the cytoplasm of the infected bat cells which are hallmarks of active viral replication (Figure 1C). This represents the establishment of the second infectious isolate of Lloviu cuevavirus.

## Discussion

Lloviu virus has now been detected in three distant countries, suggesting a more widespread occurrence of the virus than previously anticipated (Figure 1A) [1-4]. Schreibers’ bats are still the only identified host for the virus, and it is conceivable that LLOV might be present throughout the entire geographic range of this bat species.

Considering the zoonotic potential of LLOV, there is an urgent need to better understand the evolution, ecological background, and molecular features of LLOV. Our optimised qRT-PCR system combined with a novel primer set allows for the rapid and sensitive screening for LLOV viral RNA in various samples derived from bats. Large-scale screening of archived and newly collected samples would help to investigate our hypothesis about the wide geographic distribution of the virus among these bats.

The high viral load in the blood sample of the single LLOV-positive bat is in accordance with recent reports on LLOV in Schreibers’ bent-winged bats in Hungary and with previous observations for other filoviruses in bats [10,11]. LLOV-positive bats were detected in September both in Hungary and in Italy, although it is not yet clear if this points to a potential seasonality of the virus or if this reflects a random occurrence pattern. In case of European Bat Lyssavirus type 2, the peak season of virus prevalence correlated with the autumn swarming season of the bats [12].

The most important achievement of this article is the establishment of an additional infectious LLOV isolate from a bat sample using the SuBK12-08 cells, demonstrating that this cell line is highly susceptible to LLOV infection [3]. This is consistent with a previous report which found these cells to be highly susceptible to VSV pseudotyped with LLOV GP [13].

Notably, there is still no evidence for actual spillover of LLOV from bats to humans, but the repeated isolation of infectious LLOV from Schreibers’ bent-winged bats highlights the need for additional studies to better understand potential spillover mechanisms. Urine and faeces of LLOV-infected bats were found to be negative in a recent study [3]. Extensive surveillance and sequencing efforts are essential to understand the potential role of Schreibers’ bats in the natural circulation of this virus and its evolution. Of note, these bats are mainly considered to be cave-dwelling, although in rare cases their ecology can adapt in response to habitat disruption. Indeed, the increasing discovery in Italy of roosts hosting hundreds of Schreibers’ bent-winged bats within urban settings raises concerns about the increasing risk of both direct bat-human contact as well as indirect contact through the spillover to domestic animals [14]. This highlights the importance of the conservation of bats and their habitats as a means for limiting potential zoonotic spillovers to humans [15].

Cases of mass die-offs of Schreibers’ bent-winged bats have been reported across Europe, including areas where LLOV was detected [6]. This report describes the detection of LLOV from live-sampled animals that were apparently healthy, with no reported unusual mortality events among these bats in the area. Investigating the circulation of LLOV among populations of this bat species, including the estimation and evaluation of mortality events, is crucial not only for public health but also for the conservation of Schreibers’ bent-winged bats that are classified as Vulnerable on the IUCN Red List.

## Acknowledgements

The authors would like to express their sincere thanks to Andrea Lombardo, Pamela Priori, Francesca Festa, Martina Castellan and Petra Drzewnioková for their support in the collection of samples.

## Supplemental Online Material

