## Supplementary materials for "Isolation and genome characterization of Lloviu virus from Italian Schreibers’ bent-winged bats"

**Supplementary material**

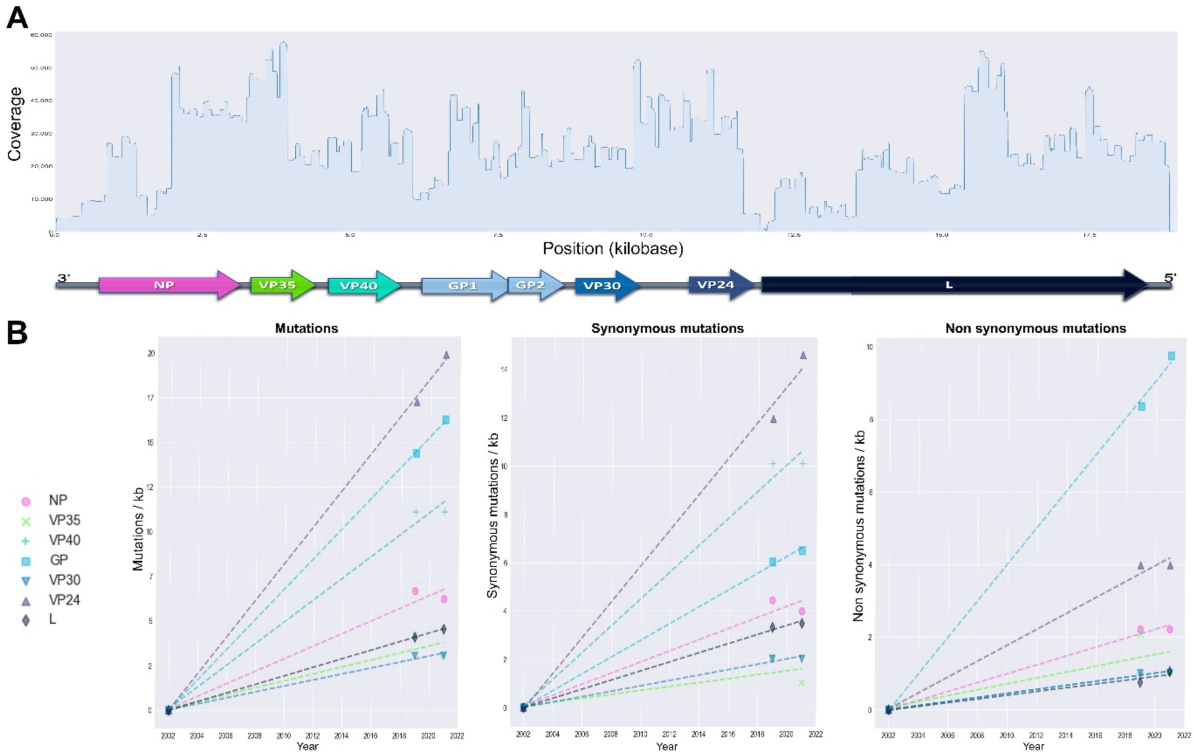


**Figure S1. (A)** The plot shows the coverage data of the Italian LLOV isolate sequencing results (ON186772) with a schematic genome representation below. **B,** The graphs show the number of mutations normalized to gene length for the 7 LLOV genes.

**Table S1.** Non-synonymous mutations comparing the three currently available complete coding sequences of Lloviu virus. Reference: NC_016144.1, Hungarian: MW775011, Italian: ON186772.

| Gene | Nucleotide position | aa position | Spanish aa position | Hungarian aa position | Italian aa position | Hungarian substitution per kb | Hungarian dN/dS ratio | Italian substitution per kb | Italian dN/dS ratio |
|---|---|---|---|---|---|---|---|---|---|
| NP | 1229 | 410 | N | T | T | 2,22 | 0,50 | 2,22 | 0,56 |
| NP | 1336 | 446 | S | P | P | | | | |
| NP | 1859 | 620 | N | S | S | | | | |
| NP | 2005 | 669 | E | K | K | | | | |
| NP | 2111 | 704 | I | T | T | | | | |
| VP35 | 487 | 163 | A | T | A | 2,08 | 1,00 | 1,04 | 1,00 |
| VP35 | 622 | 208 | L | F | F | | | | |
| VP40 | 101 | 34 | P | L | L | 1,01 | 0,10 | 1,01 | 0,10 |
| GP | 493 | 165 | H | Y | Y | 8,36 | 1,38 | 9,75 | 1,50 |
| GP | 1071 | 358 | S | P | P | | | | |
| GP | 1095 | 366 | A | T | T | | | | |
| GP | 1249 | 417 | S | N | N | | | | |
| GP | 1269 | 424 | E | K | K | | | | |
| GP | 1333 | 445 | L | P | P | | | | |
| GP | 1339 | 447 | L | P | P | | | | |
| GP | 1383 | 462 | S | P | P | | | | |
| GP | 1393 | 465 | V | A | A | | | | |
| GP | 1394 | 465 | V | A | A | | | | |
| GP | 1399 | 467 | Y | H | H | | | | |
| GP | 1431 | 478 | T | P | P | | | | |
| GP | 1439 | 480 | L | P | P | | | | |
| GP | 1446 | 483 | P | S | S | | | | |
| GP | 1462 | 488 | I | I | T | | | | |
| GP | 1463 | 488 | I | I | T | | | | |
| GP | 1476 | 493 | S | P | P | | | | |
| GP | 1537 | 513 | V | A | A | | | | |
| GP | 1584 | 529 | H | H | Y | | | | |
| GP | 2058 | 687 | T | A | A | | | | |
| GP | 2076 | 693 | I | V | V | | | | |
| VP30 | 583 | 195 | A | T | T | 1,01 | 0,50 | 1,01 | 0,50 |
| VP24 | 46 | 16 | F | L | L | 3,98 | 0,33 | 3,98 | 0,27 |
| VP24 | 472 | 158 | D | N | N | | | | |
| VP24 | 554 | 185 | C | Y | Y | | | | |
| L | 237 | 79 | I | I | M | 0,76 | 0,23 | 1,06 | 0,30 |
| L | 3292 | 1098 | G | R | R | | | | |
| L | 3467 | 1156 | D | G | G | | | | |
| L | 4874 | 1625 | R | R | K | | | | |
| L | 5159 | 1720 | S | L | L | | | | |
| L | 5171 | 1724 | R | Q | Q | | | | |
| L | 5239 | 1747 | T | S | S | | | | |
